# Behavioral Imitation with Artificial Neural Networks Leads to Personalized Models of Brain Dynamics During Videogame Play

**DOI:** 10.1101/2023.10.28.564546

**Authors:** Anirudha Kemtur, Francois Paugam, Basile Pinsard, Yann Harel, Pravish Sainath, Maximilien Le Clei, Julie Boyle, Karim Jerbi, Pierre Bellec

## Abstract

Videogames provide a promising framework to understand brain activity in a rich, engaging, and active environment, in contrast to mostly passive tasks currently dominating the field, such as image viewing. Analyzing videogames neuroimaging data is however challenging, and relies on time-intensive manual annotations of game events, based on somewhat arbitrary rules. Here, we introduce an innovative approach using Artificial Neural networks (ANN) and brain encoding techniques to generate activation maps associated with videogame behaviour using functional magnetic resonance imaging (fMRI). As individual behavior is highly variable across subjects in complex environments, we hypothesized that ANNs need to account for subject-specific behavior to properly capture brain dynamics. In this study, we used data collected while subjects played Shinobi III: Return of the Ninja Master (Sega, 1993), an action-platformer videogame. Using imitation learning, we trained an ANN to play the game while closely replicating the unique gameplay style of individual participants. We found that hidden layers of our imitation learning model successfully encoded task-relevant neural representations, and predicted individual brain dynamics with higher accuracy than models trained on other subjects’ gameplay. Individual-specific models also outperformed a number of baselines to predict brain activity, such as pixel inputs, or button presses. The highest correlations between layer activations and brain signals were observed in biologically plausible brain areas, i.e. somatosensory, attention, and visual networks. Our results demonstrate that training subject-specific ANNs can successfully uncover brain correlates of complex behaviour. This new method combining imitation learning, brain imaging, and videogames opens new research avenues to study decision-making and psychomotor task solving in naturalistic and complex environments.

## Introduction

Neuroimaging studies of human cognition increasingly rely on naturalistic paradigms that employ dynamic, rich and complex stimuli (***Sonkusare et al., 2019***). In most of the previous works the tasks studied and stimuli used have been mostly passive, such as watching videos (***Vanderwal et al., 2019***; ***Eickhoff et al., 2020***), or free reading (***Hsu et al., 2019***). However, these tasks are still far from the interactive environment the brain normally operates in. Videogames have been shown to strongly engage participants’ attention, emotions, motor and decision-making abilities (***Anderson et al., 2011***; ***Bavelier et al., 2012***; ***Palaus et al., 2017***). As such, videogames offer a powerful framework to model the integration of brain dynamics across the perception-to-action loop (***Bellec and Boyle, 2019***). Analyzing neuroimaging data collected during videogame play is however challenging, as mapping game events with cognition often requires some apriori choice of annotations which are usually generated manually (***Mathiak and Weber, 2006***; ***Weber et al., 2006***). In this work we explore a new method to characterize brain activity during videogame that leverages the full extent of the games complexity by using artificial neural networks (ANNs) and brain encoding (***Naselaris et al., 2011***).

Brain encoding has emerged as an alternative method to traditional linear general models based on manual annotations for producing brain maps. Brain encoding is especially attractive when stimuli are rich and it is not obvious what stimuli features may be driving brain activity. In a brain encoding analysis, a computational model is first trained to do a task of interest using a rich sensory data stream as input (e.g. annotation of objects from pixel-level image data), and the same model is then used to predict the brain responses while performing the same task (***Schrimpf et al., 2018***). Many brain encoding studies have successfully applied deep neural networks to model the functioning of different brain functional systems, such as vision (***Yamins and DiCarlo, 2016***; ***Hong et al., 2016***), audition (***Kell et al., 2018***), and language (***Caucheteux and King, 2022***), leading to a better understanding of the neural basis of these cognitive functions. However, these previous applications of ANNs have focused mainly on passive sensory processing and much less on active tasks involving decision-making.

A recent work demonstrated that ANNs can successfully be applied to brain encoding in videogames, specifically for classic Atari games (***Cross et al., 2021***). However, Cross and colleagues relied on a generic reinforcement learning model whose behavior was not specifically tuned to the subjects’ gameplay. Although this generic model approach may have been adequate for relatively simple games, it does not leave much room for exploring inter-individual behavioral variations during videogame play. Of particular interest is the fact that individual human videogame play can be rich and diverse given a sufficiently complex environment (***Duarte et al., 2020***; ***Bavelier and Green, 2019***).

In this work, we aimed to propose a new method to encode brain activity with ANNs during videogame play. Our central hypothesis was that individual behavior is key to training good ANN models of brain activity for complex videogame environments. Specifically, we tested that ANNs trained to closely imitate individual gameplay in the action game Shinobi III would translate to a more accurate brain encoding model of that individual than models imitating the behaviour of other individuals (see Figure 1 for an outline of the approach). We also compared individual-specific ANNs with a number of baselines, including annotation of button presses (which capture fine-grained motor behavior) and brain encoding using vision inputs (which capture fine-grained visual stimuli). This study is thus designed to establish behavioral imitation using ANNs as an automated and effective method to produce brain maps for videogames.

**Figure 1.**
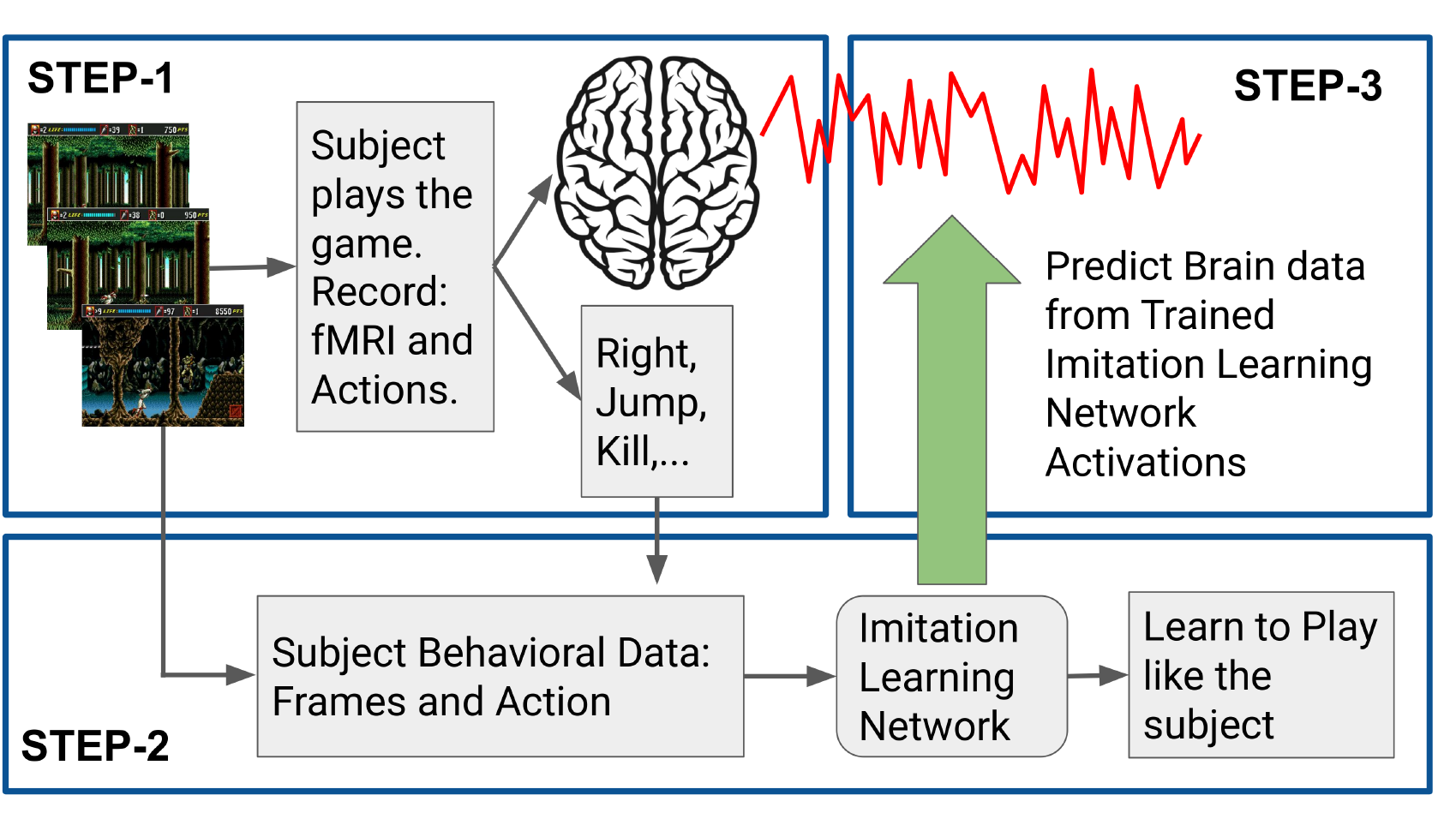
Experiment Pipeline. **Step-1**: Record videogame frames, fMRI data, and key presses of the subject. **Step-2**: Train an imitation learning neural network using frames and action data, such that it learns to play with a gameplay style similar to the subject. **Step-3**: Use activations from the neural network model to linearly predict the fMRI data.

## Materials and Methods

The summary of our experiment pipeline is shown in Figure 1. In the sections below, we describe in detail, the dataset and its collection (STEP-1 in Figure 1), followed by a description of the Imitation learning neural network and its training (STEP-2) and then how the brain encoding was performed (STEP-3).

### Dataset

The shinobi dataset used in this project was obtained from the Courtois-Neuromod databank (***Boyle et al., 2020***). The shinobi dataset consists of both fMRI and behavioral data acquired while participants played the videogame Shinobi III: Return of the Ninja Master (Sega, 1993). Data was collected on four right-handed subjects (2 women) aged from 33 to 49 years old during acquisition.

#### Shinobi videogame

In this task, the subjects had to play Shinobi III: Return of the Ninja Master, an action-scrolling platformer in which the player incarnates a ninja trying to free their village from a fictitious group of mercenaries. We selected three levels of the game for their gameplay similarity; levels 1, 4, and 5. In these levels, a player starts on the far left of the map and has to reach and complete a boss-fight located at the far right of the map. To do so, a player can move, crouch, jump, and hit enemies while collecting bonuses and avoiding various obstacles. The full completion of a level usually takes about 2 to 3 minutes, and a repetition of the level was considered complete when a) a player successfully beat the boss of that level, or b) a player dies.

The game was emulated using OpenAI’s gym-retro (v0.17) (***Nichol et al., 2018***), a toolbox that allowed the extraction of game frames and memory state variables, and presented using PsychoPy (***Peirce et al., 2019***).

#### Experiment setup

In phase one of the experiment participants were first instructed to play outside of the MRI scanner our version of the Shinobi video game using a computer monitor or TV. Subjects played until they reached a proficient level of skill, determined using performance metrics such as maximum score reached and speed of level completion (***Harel et al., 2023***).

Once subjects reached the proficient level of play they were invited to our MRI facility to play more sessions of the shinobi task for phase two. We collected behavioral data (videogame frames and controller key presses) and fMRI BOLD activation data as they played in the scanner. Around 10 sessions per subject were recorded with each session lasting for about 40 minutes and composed of multiple runs. One run of the Shinobi task comprises sequential repetitions of levels 1, 4, and 5 (in this order) looping back to level 1 when the first three levels are completed. The runs lasted for at least 10 minutes and ended when the current repetition ended after the 10 minutes had elapsed.

For this project, only Phase Two data (behavioral and fMRI) was used. Few runs and sessions were not usable based on quality checks and hence were excluded.

The data used per subject is shown in Table 1.

**Table 1.** dataset details.

| Subject | Total sessions | Total runs | Data (in hours) |
| --- | --- | --- | --- |
| 01 | 12 | 53 | 7.20 |
| 02 | 8 | 37 | 5.41 |
| 04 | 8 | 40 | 6.02 |
| 06 | 10 | 43 | 6.35 |

#### fMRI data

MRI data was collected using a 3T Siemens Prisma Fit MR scanner, a 64-channel head coil. fMRI data was collected using an accelerated simultaneous multi-slice imaging sequence (***Setsompop et al., 2012***) with a spatial resolution of 2mm isotropic, and a Repetition Time (TR) of 1.49s. Visual stimuli were projected onto a screen via a waveguide and presented to participants on a mirror mounted on the head coil. For a more detailed description of the MRI or fMRI sequences and set up are described at Courtois Neuromod’s Project documentation page: https://docs.cneuromod.ca.

#### Data Preprocessing

*Behavioral data* was recorded at 60Hz. However, it was observed that humans play the game at about 10 to 15Hz (min press duration of 4-6 samples), which translated to each controller key press being repeated 4 to 6 times in our recordings. Hence, we downsampled the behavioral data to 12Hz. This resulted in having 18 videogame frames + controller key presses for every 1.49 seconds (Duration of each fMRI TR). Videogame frames were converted from RGB to grayscale images. Further, we also cropped out the top portion of the videogame frame where the game score/health/ammunitions are displayed.

*fMRI data* was preprocessed using the fMRIprep pipeline (***Esteban et al., 2019***), using the “long term support” version 20.2.3. A high pass filter of 0.01 Hertz and Spatial smoothing of fwhm = 8mm were applied to the data and [motion + wm_csf + global] confounds were then regressed-out using nilearn’s load_confounds package. Finally, voxel space fMRI data was converted into parcel space using MIST-444 atlas (***Urchs et al., 2019***).

### Imitation learning

Imitation learning can be approached as human observational learning, having an expert perform the desired behavior and make our policy imitate the expert. This would fall under the category of Reinforcement learning algorithms known as Imitation learning. In this study, we want to see if training a network to play with the gameplay style of the subject better encodes the subject’s brain data, and we use imitation learning to train our behavior encoding model.

#### Model architecture

We use a Deep neural network-based imitation learning model. The network architecture of our model is shown in Figure 2. As the objective here is to analyze a video game frame and determine the appropriate action to mimic the subject’s behavior, the model must process visual data and maintain a record of actions taken in the preceding timestep. Therefore, our imitation learning model consists of four stacked Convolutional layers, followed by four Long Short-Term Memory (LSTM) layers. The outputs from the LSTM are then fed into a Fully Connected (FC) layer, which computes softmax probabilities over the action space. There are 2^7^ = 128 possible actions from all possible combinations of 7 button presses.

**Figure 2.**
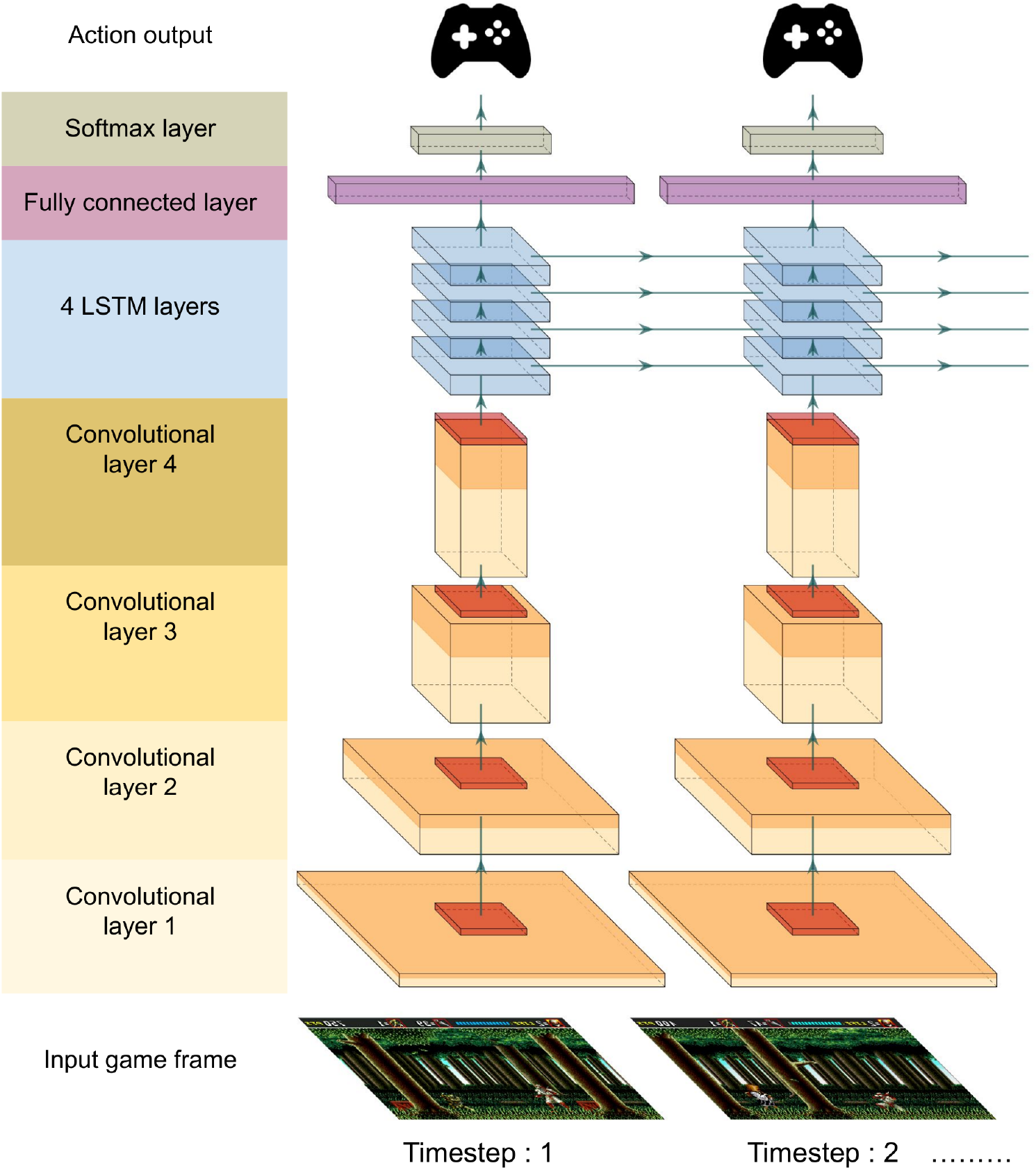
Model Architecture is composed of four Convolutional layers, four LSTM layers, fully connected layer to output softmax probabilities over the action space.

The architecture of our imitation model is shown in Figure 2 and layer details are described in Table 2.

**Table 2.**
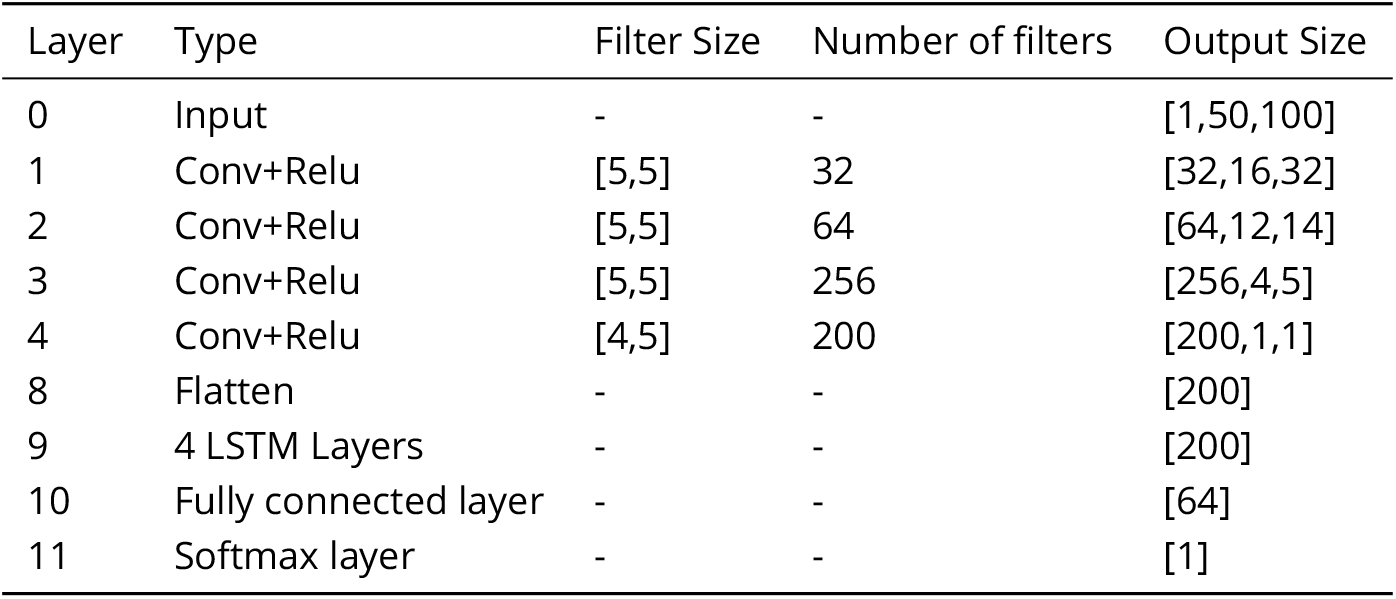
Architecture details.

#### Training

##### Behaviour Cloning

The simplest and one of the earliest approaches to IL is Behaviour Cloning (BC). BC aims to learn the expert’s policy by treating it as a supervised learning problem. Given the expert state-action pairs, treated as i.i.d examples, we apply supervised learning by minimizing the loss between the model’s predicted action and the subject’s recorded action for that game frame.

##### Loss function

There is a prominent class imbalance in the action sequences, notably the “go-right” action is the most frequent step (due to the nature of a right-scroller game). A gameplay trajectory requires every action in a particular sequence, and removing some actions to solve the class imbal-ance problem would result in an entirely different trajectory. We solved this class imbalance with a weighted negative log-likelihood loss function (based on the frequency of each action in the data) so that errors on frequent actions are less penalized than infrequent actions ones. This allows the model to focus more on the infrequent actions while maintaining the original game trajectory.

##### Implementation

The model was implemented using PyTorch 1.6 in Python 3.6. The loss was then backpropagated through the model and weights were updated using the Adam optimizer (***Kingma and Ba, 2019***). Early stopping (***Zhang et al., 2005***) was employed to prevent overfitting and the best model was stored for validation. Hyperparameter selection is important to train a performant deep-learning model. We carried out an exhaustive search for the optimal hyperparameters following well-accepted principles and approaches in the field (***Smith, 2018***). The hyperparameter values found are provided in Table 3.

**Table 3.** Training Hyperparameters.

| Hyperparameter | Value |
| --- | --- |
| LSTM sequence length | 18 |
| Batch size | 50 |
| Training epochs | 3000 |
| Learning rate | 0.00005 |

### Brain encoding

To analyze the relationship between the feature space learned by the ANN and the subject’s brain data (***Wang et al., 2015***; ***Mohsenzadeh et al., 2019***), we performed a brain encoding approach using linear mapping.

The details of this analysis are summarised in Figure 3. First, we reduced the dimensions of the activations as we have limited data points and hence cannot use very large feature vector dimensions to train the linear regression model. Activations from the four CNN layers, LSTM layer, and FC layer are reduced in dimension (output vector dimension - 800) using a principal component analysis (PCA). Second, to get encoding features at the fMRI sampling rate, we applied another PCA (output vector dimension - 300) to compress information from 1TR=1.49-second windows, independently for each layer. Finally, the PCA vectors resulting from the second PCA from each layer are concatenated to compose the final feature space vector used for ridge regression. To account for the hemodynamic delay, the final feature space vector of the imitation model at time t is mapped to fMRI data at time t+X seconds, where the best encoding was found to be at X=6, consistent with established properties of the hemodynamic response function (***Glover, 1999***). A separate ridge regression model is fit for each parcel.

**Figure 3.**
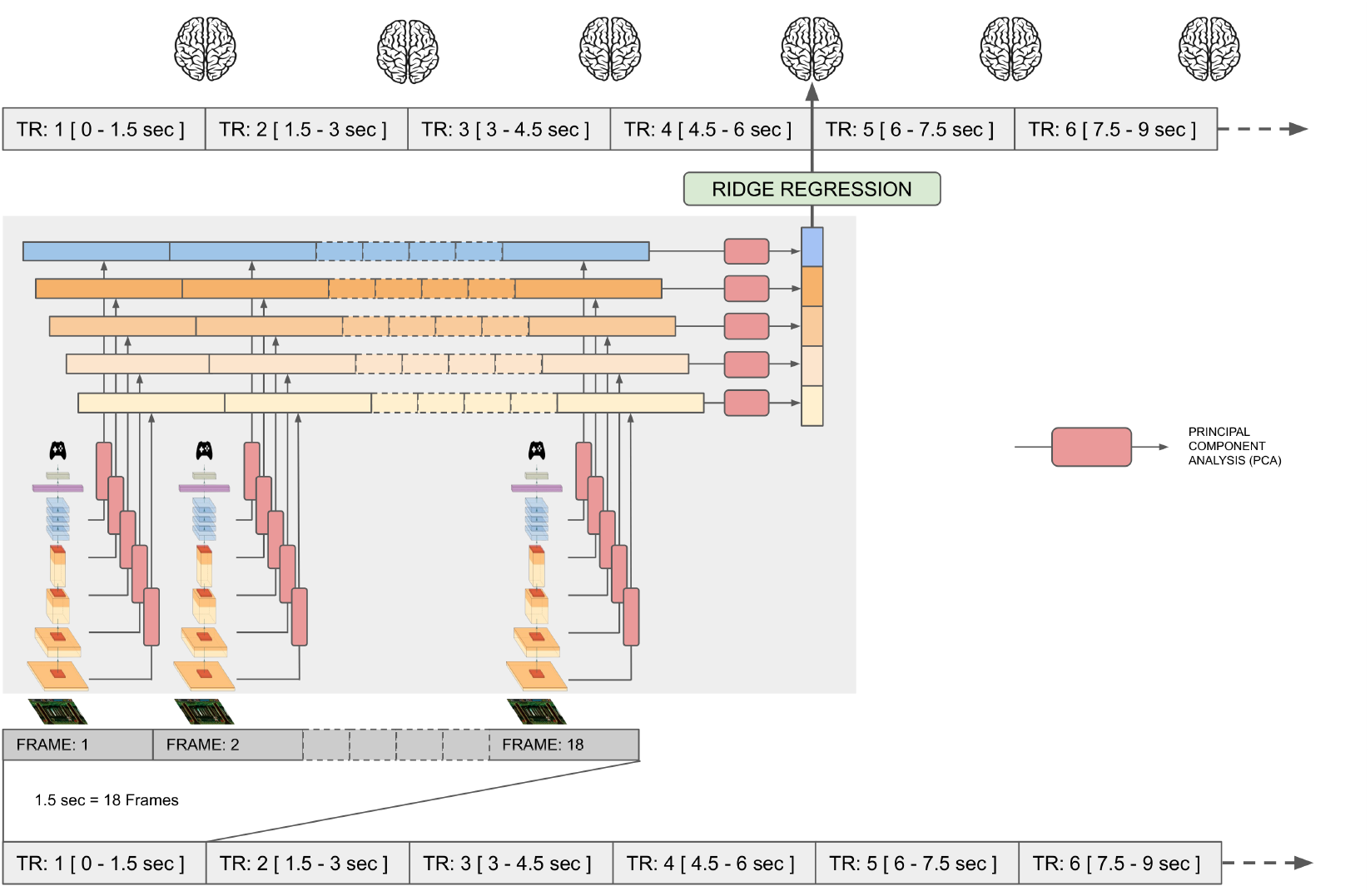
Brain encoding framework. The subject’s gameplay frames are passed into a trained imitation learning model and its internal network activations are reduced in dimension using PCA. These resulting features are used to predict fMRI BOLD responses using ridge regression.

### Control models

To test whether training an ANN to mimic the behavior of a given individual endows it with higher brain encoding performance in that individual, we compared brain encoding performance across a range of control models described below.

#### Untrained model

When pixel input is processed by an untrained model, the inherent structure of Convolutional and LSTM layers enables the capture of useful features. To assess how much variance is captured solely due to the model’s architecture, we use an untrained model with randomly initialized weights as one of our control models.

#### Pixel space PCA model

To capture the amount of brain data variance explained just by the videogame frames, we applied a PCA on the pixel RGB input (output vector dimension - 1000) and used it to train the ridge regression model.

#### Action space model

We use the controller key presses of the subject to predict the brain data. This allows us to capture the amount of brain data variance explained just by the final action space.

## Results

### ANNs trained to imitate individual gameplay successfully encoded brain activity

#### Behavioral results

We first aimed to train ANNs that could imitate the behavioral gameplay of individual subjects. We trained a deep ANN composed of vision (CNN) and recurrent (LSTM) layers using behavioral cloning, to predict button presses directly from pixel-level video frames from the game (out of 128 possible actions)(Figure 2). Leave-one-session-out cross-validation was performed, and the model imitated the subjects’ actions with an accuracy of 62-68%; sub-01: 62%, sub-02: 64%, sub-04: 68%, sub-06: 66% (Chance level: 0.8%). To complement these quantitative performance metrics, we also used trained models to play the game for entire levels, by feeding the actions predicted by the model back into the game emulator. Qualitatively we found that trained models performed adequately, being able to progress through levels and often complete them, and taking sequences of actions analogous to individual subjects’ gameplay. A demo video of an imitation learning model playing the game with a subject-specific style is available at this [Link](https://figshare.com/articles/media/imitation_learning_demo_mp4/24667863).

#### Brain encoding

Next, we tested whether the representations learned by individual imitation models could be used to predict task-related fluctuations in brain activity, for the same subjects. We implemented a brain encoding analysis, where the video frames of the game of a player were fed into the imitation model. The resulting model layers activations were used as input to a Ridge regression that predicted brain activity in all of the brain parcels (Figure 3).

The coefficient of determination *R*^2^ was calculated between the predicted fMRI data from the ridge regression model and the actual fMRI data for each parcel (Figure 4). We observe that the imitation learning model predicted brain dynamics with high accuracy in distributed brain regions. Both the values of brain encoding values and the spatial distribution of regions with the highest encoding scores were largely consistent across subjects. Among these, the visual cortex (in particular the dorsal stream), sensorimotor cortex, as well as some frontoparietal areas showed consistent higher prediction, similar to the results obtained by Cross and colleagues (***Cross et al., 2020***).

**Figure 4.**
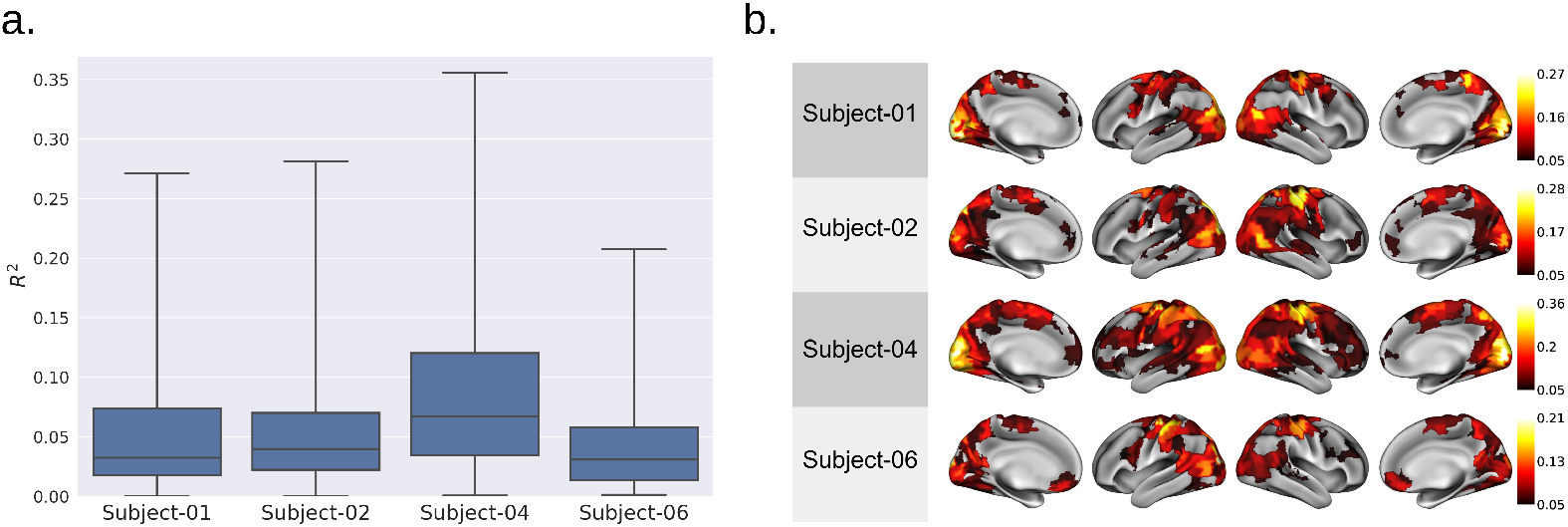
Percentage of variance explained in actual fMRI BOLD data by responses predicted by the model. The fMRI data were predicted from the internal representations of separate imitation learning models, each trained on the corresponding subject’s gameplay. Panel **a**: Distribution of brain encoding *R*^2^ (percentage of variance explained in actual fMRI BOLD data by responses predicted by the model) shown across four subjects. For each subject distribution of *R*^2^ across 444 parcels is plotted using a box plot showing the minimum, first quartile, median, third quartile, and maximum. Panel **b**: *R*^2^ across four subjects plotted on a brain map.

#### Layer wise analysis

In this section, we analyze what effect the early layers vs later layers of the model have on brain encoding and which brain regions each of them help to encode better. Apart from the encoding model that uses all the layer features we consider two additional models. Early layer model, where only the initial two layers (Convolutional layer-1, Convolutional layer-2) features were used, and the last layers model where only the last two layers (Convolutional layer-4 and LSTM layer) features were used to predict the brain data using ridge regression.

The *R*^2^ values of these 2 models were projected onto the individual brain, as shown in Figure 5. We found that the early layer better predicted the early visual cortex activation (shown in Figure 5 by rows (B) and (A-C)), while the last layers of the model better predicted the higher visual cortex, motor, and somatosensory regions (shown in the Figure 5 by rows (C) and (A-B)).

**Figure 5.**
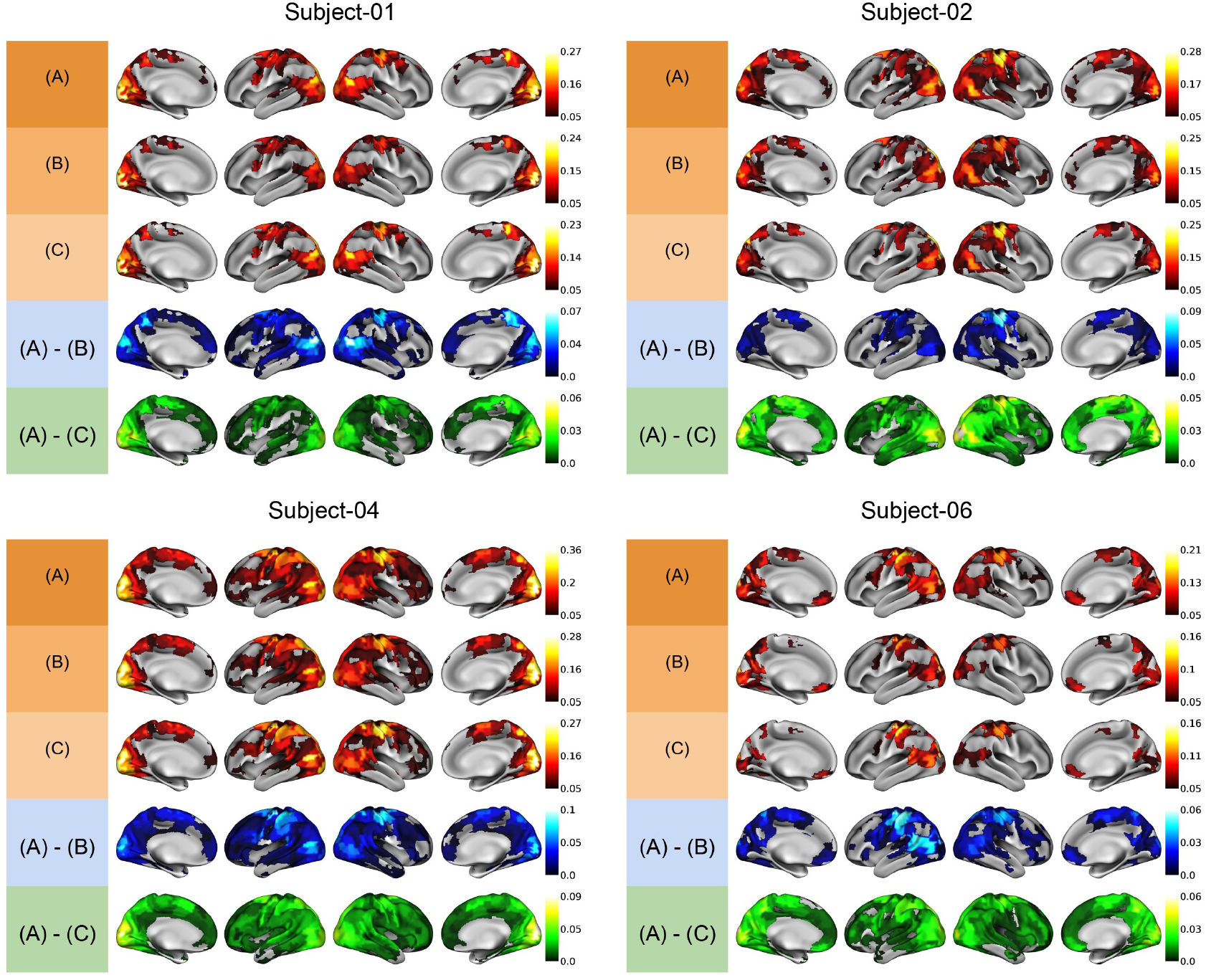
*R*^2^ values on a brain map shown for each of the four subjects. Comparison of encoding based on which layers from the trained model were used. **A**: All the layers features were used. **B**: Only early two convolutional layer features were used. **C**: The last convolutional layer and LSTM layer features were used. **A-B**: *R*^2^(All layers) - *R*^2^(Early layers). This shows the effect of the latter layers in the model. **A-C**: *R*^2^(All layers) - *R*^2^(Last layers). This shows the effect of early layers in the model.

### Subject-specific imitation translates into better brain encoding than models trained to imitate other subjects’ gameplay

#### Behavioral results

We saw that the imitation learning model was able to predict behavioural actions with an accuracy of up to 65%. Here, we tested whether these imitation models were subject-specific. Specifically, we evaluated if the imitation learning model trained on a subject’s gameplay data can model the behavior of that same subject better than a model that knows how to play the game but in a different gameplay style, drawn from the other three subjects. The results showing the within and between subject comparison are shown in Figure 6a. While the imitation learning model trained on the subject’s own gameplay was able to replicate the gameplay data up to 65% accuracy, the models trained on a different gameplay style only achieved 35% accuracy. This result was observed consistently across all four subjects.

**Figure 6.**
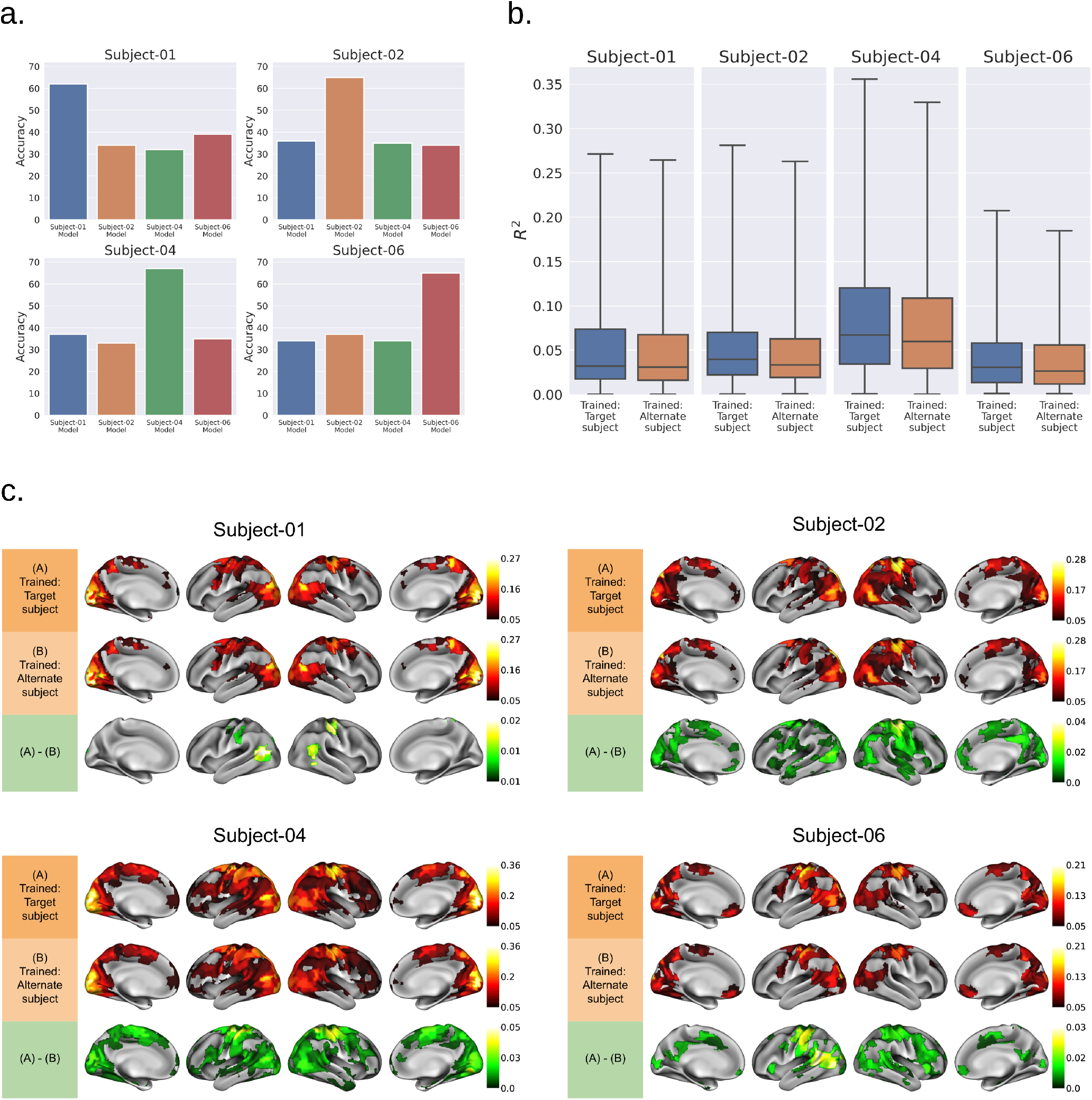
Panel **a**: Behavioral imitation learning accuracy across four subjects. X-axis showing which subject the model was trained on. Panel **b**: Distribution of brain *R*^2^ comparing encoding performed using model trained on target(same) subject vs alternate(best out of rest three) subject. Panel **c**: *R*^2^ values on a brain map shown for each of the four subjects. **First row:** Brain map of *R*^2^ from the trained model (same subject gameplay). **Second row:** Brain map of *R*^2^ from the Trained model (Different gameplay). **Third row:** Brain map of *R*^2^ (trained model: same subject gameplay) - *R*^2^ (trained model: different gameplay), where the *R*^2^ values are subtracted parcel-wise and plotted (only parcels where the difference is significant at q<0.05 are colored).

#### Brain encoding

Next, we wanted to see if the subject-specificity of imitation models translated into better brain encoding (higher *R*^2^). Specifically, we tested if a subject-specific imitation model was able to encode the brain data of that target individual better than imitation learning models trained on a different, alternative subject gameplay. This alternative subject was selected as the one most behaviorally similar to the target, i.e. whose behavioral model reached the highest accuracy on the target subject (sub-01 was paired with sub-06, sub-02 with sub-01, sub-03 with sub-01 and sub-04 with sub-02). We passed prerecorded gameplay frames of the target subject into these two models: (1) the imitation model trained on the target subject and (2) the imitation model trained on the best behavioural alternative subject. The internal activations from these models were used to predict brain data of the target subject using ridge regression.

Distribution of *R*^2^ across the parcels for each subject-specific model and the best behavioural alternative model were plotted (Figure 6b). Across all four subjects, we observed that the target (same-subject) imitation model was better at encoding the brain data than the alternative model selected for that subject. These differences were significant at *q* < 0.05 (Wilcoxon signed-rank test with false discovery rate correction for multiple comparisons across parcels). Same-subject imitation learning models better encode brain data than their corresponding alternative subject model in 6%, 35%, 37%, 26% of parcels in Subjects 01, 02, 04 and 06 respectively.

Figure 6c shows in which brain regions the subject-specific model encoded the brain dynamics better than the best behavioural alternate subject model. Specifically, we found improved encoding in motor, somatosensory, and visual cortices, compared to models trained with different gameplay. These areas are key to processing the input of the task (visual stimuli) and generating outputs (motor command).

### Subject-specific imitation models outperformed a series of control models for brain encoding

We compared brain encoding with subject-specific imitation models to a series of control models (1) PCA features derived from gameplay input pixel (Pixel-space model); (2) one-hot encoding of the controller button presses (Action-space model); and, (3) passing the subjects’ gameplay frames to a CNN-LSTM model with random weights (Untrained model).

Distribution of *R*^2^ across the parcels for imitation and control models was plotted (Figure 7). Across all four subjects, the subject-specific imitation model was better at encoding brain data than all three controls. Significant improvement in brain encoding using imitation models was found in about 40% of parcels (average across subjects) when compared to the Untrained model, and in about 85% of parcels (average across subjects) when compared with the Pixel-space and Actionspace control models. Brain maps showing which regions were better encoded by the imitation models than the controls can be found in Figure 8.

**Figure 7.**
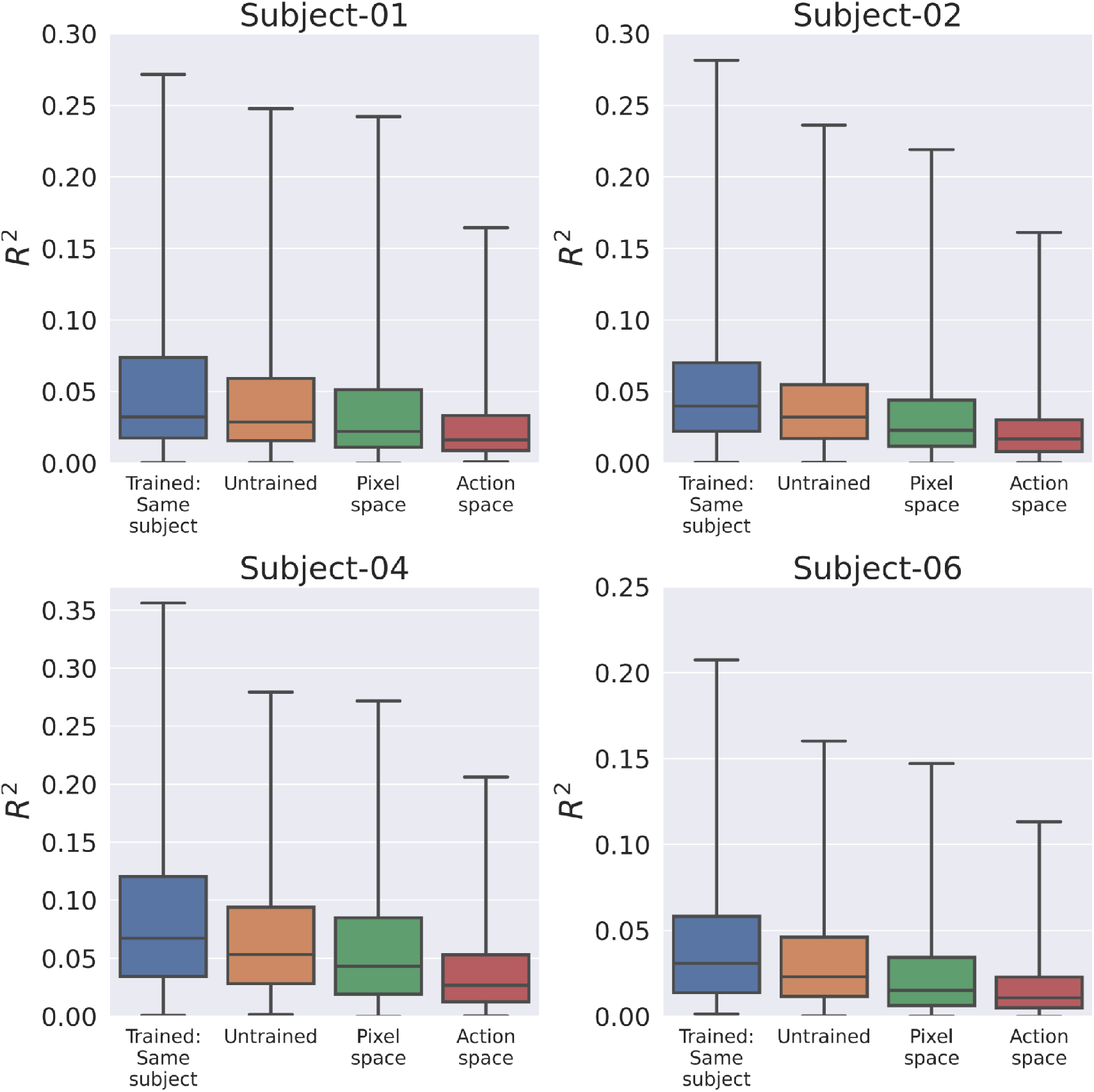
Distribution of brain *R*^2^ comparing encoding performed using model trained on target(same) subject vs three control models.

**Figure 8.**
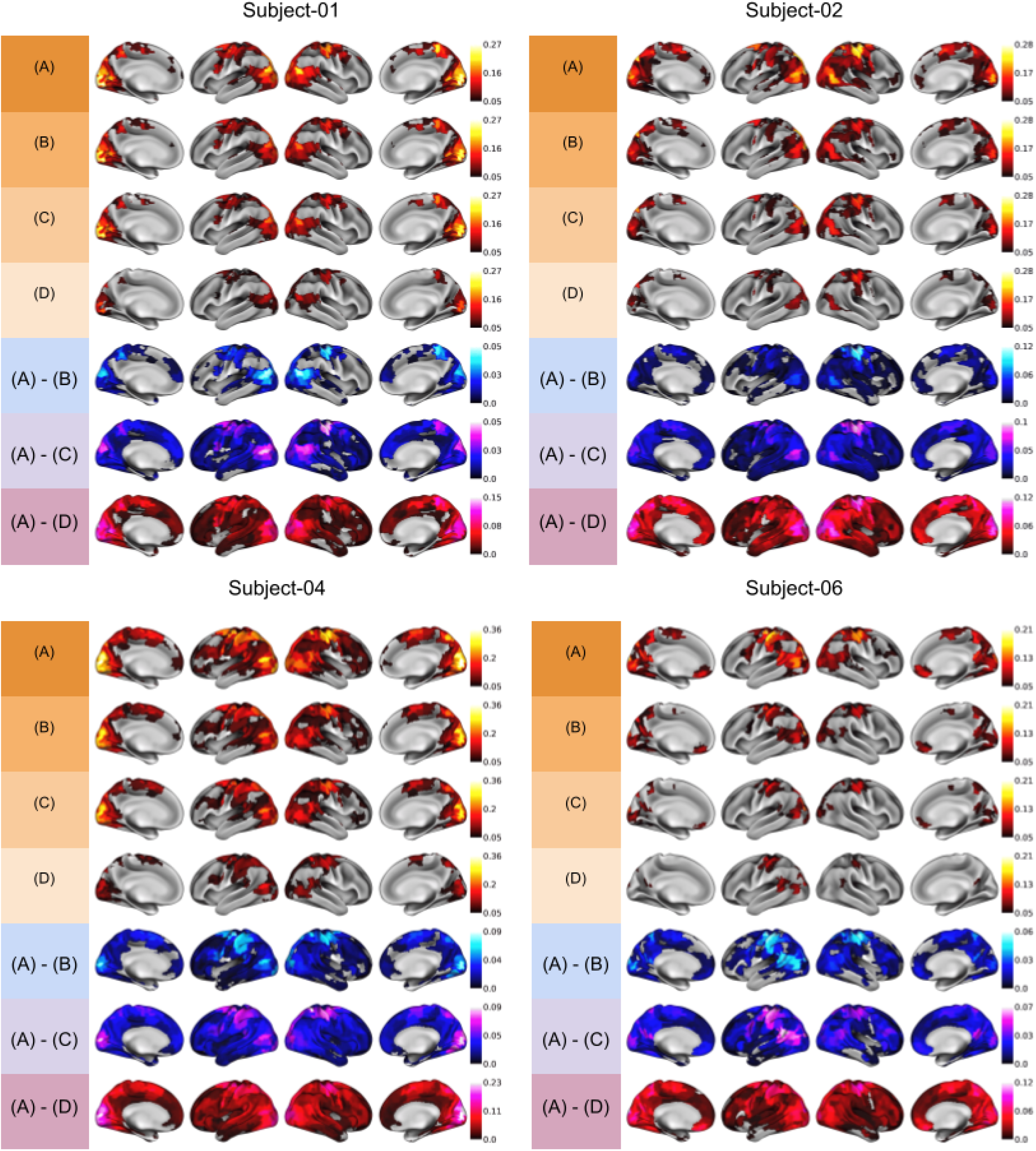
*R*^2^ values on a brain map shown for each of the four subjects. **A**: Trained model (Same subject gameplay). **B**: Untrained model. **C**: Pixel-space model. **D**: Action-space model. **A-B**: *R*^2^(Trained model) - *R*^2^(Untrained model). **A-C**: *R*^2^(Trained model) - *R*^2^(Pixel space model). **A-D**: *R*^2^(Trained model) - *R*^2^(Action space model).

## Discussion

Our individual imitation models trained with behavioural cloning show good quantitative performance (up to 70% accuracy on a hold-out test set) as well as qualitative face validity when used to play the game autonomously. It is however difficult to conclude whether these models fully imitate individual gameplay. Humans have variable behavior, and it is very likely that a player would not be able to predict their own actions 100% of the time if presented with recorded videos of their gameplay. Another critical aspect driving variations in behaviour is ongoing learning, as subjects get more proficient at the game over time. Unlike simple games like pong, with a clear objective and limited variations in strategies, the shinobi environment is rich and participants may change strategies over time, e.g. favor speed over health loss, or abandon collecting certain power ups only to revert back collecting them at a latter stage. In this experiment, subjects were extensively trained outside of the scanner prior to neuroimaging sessions in order to avoid large learning effects. However, given the complexity of the game, it is almost certain that some amount of learning and variations in strategies still happened over the imaging sessions, which cannot be modeled with our current BC approach. The study by Chen et. al. (***Chen et al., 2021***) showed when trained on various Atari games the BC approach achieves accuracies between 60% to 90% depending on the game and the size of training data. The work by Kanervisto et. al.(***Kanervisto et al., 2020***) showed that the accuracies of the BC models greatly depend on the quality of data. Hence it might not be easy to compare our accuracies to previous works which might have a difference in dataset size, quality, and nature of videogames.

We found that an ANN model trained to imitate the subject serves as a good encoding model of their own brain dynamics, with *R*^2^ values up to 35%. The encoding worked particularly well in motor, somatosensory, and visual cortices. Although these brain regions were consistent across subjects, the average and maximum *R*^2^ varied markedly across subjects. These results likely reflect systematic differences in data quality (and noise ceiling) or individual behavioral consistency across subjects. Noise in fMRI is complex, but a major source is motion (***Power et al., 2015***), and subjects had markedly different amounts of motion, consistently across scans (***Harel et al., 2023***). We also found that encoding *R*^2^ was poor in subcortical regions across all subjects, and the sequence used in our experiment had low SNR in those regions as is common with multiband accelerated fMRI. In raw image presentation, a noise ceiling can be estimated through repeated stimuli presentation to define an optimal brain response as done in this study by Allen et. al.(2022) (***Allen et al., 2022***), which was not possible to implement with videogame whose gameplay varied across repetitions and subjects. The only previous study of brain encoding with ANNs in videogames, Cross et. al.(***Cross et al., 2020***), also achieved maximum encoding *R*^2^ up to 0.38 (average across subjects) and also had the best encoding in visual, motor, and somatosensory areas. It is not possible to directly compare *R*^2^ between the two studies however, as noise ceiling (and *R*^2^) are directly impacted by parameters such as image resolution, scanning sequence, and the amount of spatial smoothing, none of which were harmonized across the two studies.

To understand which regions of the brain were predicted better by which layers of the model, we compared brain encoding predictions from features derived from only early layers vs the last layers of the model. We found that early layers predicted the brain data better in the early visual cortex and the latter layers (which included the LSTM layer) of the model better predicted the higher visual cortex, motor, and somatosensory regions. This can be attributed to the fact that early convolutional layers capture the primitive visual features of the image similar to what happens in the early visual cortex areas, and the latter layers of the model capture the gameplay behavior, which is also exhibited in motor and somatosensory regions of the brain. These layer-wise effects on brain encoding are consistent with results observed in various brain encoding studies like Yamins et. al.(***Yamins and DiCarlo, 2016***) and Cross et. al.(***Cross et al., 2020***).

Further, we found that imitation models were subject-specific. While imitation models trained on individual gameplay were able to predict actions with an accuracy of up to 70% for this individual, imitation models trained on the gameplay of another subject were only able to predict actions accurately with up to 30-40% accuracy. This improvement in behavioral accuracy also translated to better brain encoding. Specifically, we found that individual-specific imitation models improved encoding in motor, somatosensory, and visual cortices, compared to models trained with different gameplay. These areas are key to processing the input of the task (visual stimuli) and generating outputs (motor command). While imitation models trained on different gameplay know how to play the game, their actions might be dissimilar from the target subject, as demonstrated by the loss in accuracy for predicting actions. Representations of these subject-specific actions likely explain the ability of subject-specific imitation models to capture more brain variance in motor and somatosensory areas. By contrast, brain encoding in frontoparietal cortices did not show widespread improvement. Frontoparietal cortices are involved in planning, and were expected to show improvement with subject-specific behavioral imitation models. We still note some improvements in the dorsal attentional network, which is a subset of frontoparietal regions tightly linked with motor control and attention. In the experiment setup, we asked the subjects to practice the game for about six months before they played the game inside fMRI scanner. They had reached highly consistent gameplay which might have led to high levels of automation, and little effort for planning. This may explain the limited involvement of frontoparietal networks.

Control analysis showed that neither pixel input or the actions of the player were sufficient to capture brain dynamics in a complex task like playing Shinobi. We also see that training an ANN with an imitation model is beneficial to brain encoding, and features derived from an untrained CNN-LSTM network (to which raw input was fed) did not explain any more variance than models trained on raw input game pixel data. These results were observed across all cortical regions. Some previous studies showed that the structure of a CNN network is sufficient to capture a great number of low-level image statistics prior to any learning (***Ulyanov et al., 2018***) and such untrained networks can be used to encode the brain in visual tasks (***Kim et al., 2021***).

Our results show that in complex naturalistic tasks such as videogames, the stimuli or final behavior or network architecture alone are not able to capture the brain dynamics. By contrast, a deep learning model that learns to perform the task in a way similar to the subject would be able to encode the brain dynamics of that subject better. Cross et. al.(***Cross et al., 2020***) study on videogame brain encoding, found similar results where raw statistical properties of input/output were not able to explain the variance in brain data well while playing videogames, even if their reference models was not tuned to specific individual gameplay.

One of the limitations of this study is that given the slow temporal resolution of fMRI and the rapid dynamics of video gameplay, the advantages of a subject-specific imitation learning model might not have been completely translated to better brain encoding, due to a loss of fast action dynamics. Future work using high temporal resolution imaging techniques, such as EEG or MEG, would thus complement this fMRI study.

Our proposed approach to generate brain maps in complex videogames at the individual level opens new avenues for future research. There has been a push towards using more ecologically valid tasks in the context of fMRI research, including to study brain/behavior associations, e.g. in the Healthy Brain Network project (***Alexander et al., 2017***). These ecological tasks could potentially expose some associations that would otherwise not be potentiated due to a lack of engagement from participants. The most common ecological task in fMRI is movie watching, which while more engaging than resting-state remains a fairly passive task. The prospect of using commercial videogames combined with ANNs to generate individual brain maps will enable an entire new class of paradigms to be tested for brain-behavior associations - as these maps can directly be entered in a group-level general linear model. Such active tasks may be particularly well suited for certain demographics such as children, provided the videogame is selected appropriately for the target group.

## Conclusion

In this work, we showed that a deep recurrent neural network is able to successfully imitate the actions of individual human players in a complex videogame and that the representations learned by this network can be used to perform brain encoding. Those derived imitation models were subject-specific, both in terms of behavior (button presses) and brain encoding. Subject-specific imitation models also performed significantly better than a series of controls, including direct use of button presses to predict brain activity. We found that the improvement in encoding over the control model was particularly noticeable in the task-relevant networks (motor, somatosensory, and visual cortices). This new approach could see future application in studying brain and behavior associations using brain maps generated in the context of complex, commercial videogames.

## Ethics Statement

All subjects of the CNeuromod datasets provided informed consent to participate in this study, which was approved by the ethics review board of the “CIUSSS du centre-sud-de-l’île-de-Montréal” (under number CER VN 18-19-22).

## Data and Code Availability

Data used in the paper can be requested through Cneuromod website : https://www.cneuromod.ca/. The code for the study can be found in GitHub repository [Here](https://github.com/anirudhk686/shinobi_imitation_learning).

## Author Contributions

The majority of the work described in this article was done by Anirudha Kemtur.

Dr. Basile Pinsard helped with the preprocessing of fMRI data.

Francois Paugam, Pravish Sainath, Yann Harel, Maximilien Le Clei and Julie Boyle provided useful feedback throughout the study.

Dr. Karim Jerbi and Dr. Pierre Bellec supervised the work, provided the initial idea for the project, and the data in addition to mentoring and continued feedback.

## Funding

The Courtois project on neural modeling was made possible by a generous donation from the Courtois Foundation, administered by the Fondation Institut Gériatrie Montréal at CIUSSS du Centre-Sud-de-l’île-de-Montréal and University of Montreal. The Courtois NeuroMod team is based at “Centre de Recherche de l’Institut Universitaire de Gériatrie de Montréal”, with several other institutions involved. See the cneuromod documentation for an up-to-date list of contributors (https://docs.cneuromod.ca).

AK was supported in part by a Mitacs bursary in addition to Courtois foundation grant, FP, BP, PS, MLC, and JB were entirely supported by Courtois foundation grant, PB is supported by a salary award of the “fonds de recherche du Québec - Santé” (chercheur boursier senior). KJ is supported by funding from the Canada Research Chairs program and a Discovery Grant from the Natural Sciences and Engineering Research Council of Canada.

## Declaration of Competing Interests

The authors declare that they have no known competing financial interests or personal relationships that could have appeared to influence the work.

## Notes

### Competing Interest Statement

The authors have declared no competing interest.

### Summary of Updates

Update to discussion of the article.

